# Sex Differences in Dopamine Release in Nucleus Accumbens and Dorsal Striatum Determined by Chronic Fast Scan Cyclic Voltammetry: Effects of social housing and repeated stimulation

**DOI:** 10.1101/2023.08.14.553278

**Authors:** Ivette L. Gonzalez, Christopher A. Turner, Paras R. Patel, Noah B. Leonardo, Brandon D. Luma, Julianna M. Richie, Dawen Cai, Cynthia A. Chestek, Jill B. Becker

## Abstract

We investigated sex differences in dopamine (DA) release in the nucleus accumbens (NAc) and dorsolateral striatum (DLS) using a chronic 16-channel carbon fiber electrode and fast-scan cyclic voltammetry (FSCV). Electrical stimulation (ES; 60Hz) induced DA release was recorded in the NAc of single or pair-housed male and female rats. When core (NAcC) and shell (NAcS) were recorded simultaneously, there was greater ES DA release in NAcC of pair-housed females compared with single females and males. Housing did not affect ES NAc DA release in males. In contrast, there was significantly more ES DA release from the DLS of female rats than male rats. This was true prior to and after treatment with methamphetamine. Furthermore, in castrated (CAST) males and ovariectomized (OVX) females, there were no sex differences in ES DA release from the DLS, demonstrating the hormone dependence of this sex difference. However, in the DLS of both intact and gonadectomized rats, DA reuptake was slower in females than in males. Finally, DA release following ES of the medial forebrain bundle at 60Hz was studied over four weeks. ES DA release increased over time for both CAST males and OVX females, demonstrating sensitization. Using this novel 16-channel chronic FSCV electrode, we found sex differences in the effects of social housing in the NAcS, sex differences in DA release from intact rats in DLS, sex differences in DA reuptake in DLS of intake and gonadectomized rats, and we report sensitization of ES-induced DA release in DLS *in vivo*.

**Significance Statement:** Dopamine release is not uniform or fixed. In the nucleus accumbens, pair housing, compared with individual housing, is shown to differentially affect dopamine responsiveness to stimulation in a sex-dependent and region-specific way. There are also sex differences in stimulated dopamine release in the dorsolateral striatum of intact rats, which are not seen in gonadectomized rats, indicating the hormone dependence of this sex difference. However, reuptake of dopamine was slower in females than in males, independent of gonadal hormones. Importantly, the electrical stimulation-induced dopamine release in the dorsolateral striatum of gonadectomized rats demonstrated sensitization of dopamine release *in vivo* within animals for the first time. Thus, stimulated dopamine release exhibits sex-specific neuroplasticity that is modified in females by the housing conditions.

## Introduction

There are sex differences in stimulated dopamine (DA) release *in vivo* both in the dorsolateral striatum (DLS) and nucleus accumbens (NAc) as detected with microdialysis (Castner et al., 1993; Cummings et al., 2014; Quigley et al., 2021). Furthermore, ovariectomy (OVX) of female rats attenuates, and estradiol restores DA release induced by psychomotor stimulants or K^+^ *in vivo* and *in vitro* (Becker and Ramirez, 1981; Becker, 1990; Cummings et al., 2014). When OVX females and castrated (CAST) male rats are compared using microdialysis, the cocaine-induced increase in DA is lower in OVX than in CAST in both DLS and NAc (Becker and Ramirez, 1981; Cummings et al., 2014). Furthermore, basal DA concentrations are higher in CAST than in OVX (Xiao and Becker, 1994), and stimulated increases in DA measured with microdialysis or *in vitro* vary with the estrous cycle (Becker and Ramirez, 1981; Becker and Cha, 1989).

Pair-housing has been shown to attenuate the effects of mild chronic stress in female rats but not males (Westenbroek et al., 2003). Pair-housing also attenuates motivation to self-administer cocaine in females but not in males (Westenbroek et al., 2013). Finally, in pair-housed females, the methamphetamine (METH) induced increase in DA in dialysate from the NAc is attenuated after social housing (Westenbroek et al., 2019). Thus, social housing in rats has sex-specific effects on behavior and DA function in the NAc.

In previous studies, we have repeatedly used microdialysis to measure stimulated DA increases in the NAc (Perry et al., 2015). There are significant challenges, however, with using microdialysis repeatedly. In fact, various techniques to measure DA release directly *in vivo* have struggled with spatial and/or temporal resolution (Wu et al., 2022).

Fast-scan cyclic voltammetry (FSCV) uses a carbon fiber electrode to measure oxidizable molecules, such as DA, with high temporal resolution (Garris and Wightman, 1995). FSCV has been used *in vitro* with brain slices and *in vivo* in anesthetized or freely-moving animals (Garris et al., 1997). Recordings from chronically implanted FSCV electrodes have been used to allow for studies with a more longitudinal design but introduce different challenges, such as reduced sensitivity due to immune response (Clark et al., 2010). While there have been many advancements in carbon fiber electrode technology for use in freely moving animals, there are still limitations.

In this project, we use a novel chronically implanted FSCV electrode to detect DA release in freely moving animals. This 16-channel high-density carbon fiber (HDCF) array was designed and manufactured at the University of Michigan (Huan et al., 2021). These probes maintain reliable high-yield recordings for at least two months (Welle et al., 2020). With these electrodes, we can simultaneously record from anatomically distinct subregions of the NAc, the core (NAcC), and the shell (NAcS), as well across approximately 1 mm in the DLS. With these probes, we measure basal (DA transients) and electrically stimulated (ES) DA release and reuptake in the NAcC and NAcS or the DLS in a freely moving animal reliably recorded for weeks. The location of the probe can be determined postmortem using slice-in-place technology.

In previous studies using microdialysis in the NAc, our probes were sampling from both NAcC and NAcS. With the probes used here, we investigate the effect of pair-housing on ES DA release and can now specifically determine the effects in the NAcC and NAcS in male and female rats. This allows us to determine whether we find effects of social housing and/or sex differences when NAcC and NAcS are examined specifically. We also examine whether there are sex differences in ES DA release in DLS of intact female and male rats. Finally, we investigate whether we see sex differences in ES DA release in the DLS of OVX and CAST rats and the extent to which there are changes in ES DA release in the DLS with repeated testing over four weeks.

## Materials and Methods

### Subjects

Male and female Sprague-Dawley rats were obtained from Charles River Breeding Laboratory (Portage, MI). Animals were maintained in ventilated laboratory cages on a 14:10 light/dark cycle in a temperature-controlled climate of 72 ± 2 °F. Rats had *ad libitum* access to water and phytoestrogen-free rat chow (2017 Teklad Global, 14% protein rodent maintenance diet, Harlan rat chow; Harlan Teklad). All procedures were performed according to protocols approved by the University of Michigan Institutional Animal Care and Use Committee.

#### Experiment 1

For experiments targeting the NAc, animals were approximately 40-45 days old on arrival. This experiment used 12 females and 10 males in total. Of those totals, 6 females and 5 males were individually housed. 6 of the females and 5 of the males were housed in pairs (social housing) for three weeks before surgery. One of the paired rats underwent surgery and was individually housed for 6 days for post-operative care. On day 6, pairs were reunited and remained paired until the end of the experiment. This design was implemented equally for both males and females (Figure 1). Rats were about 75 days old on their first day of testing.

**Figure 1.**
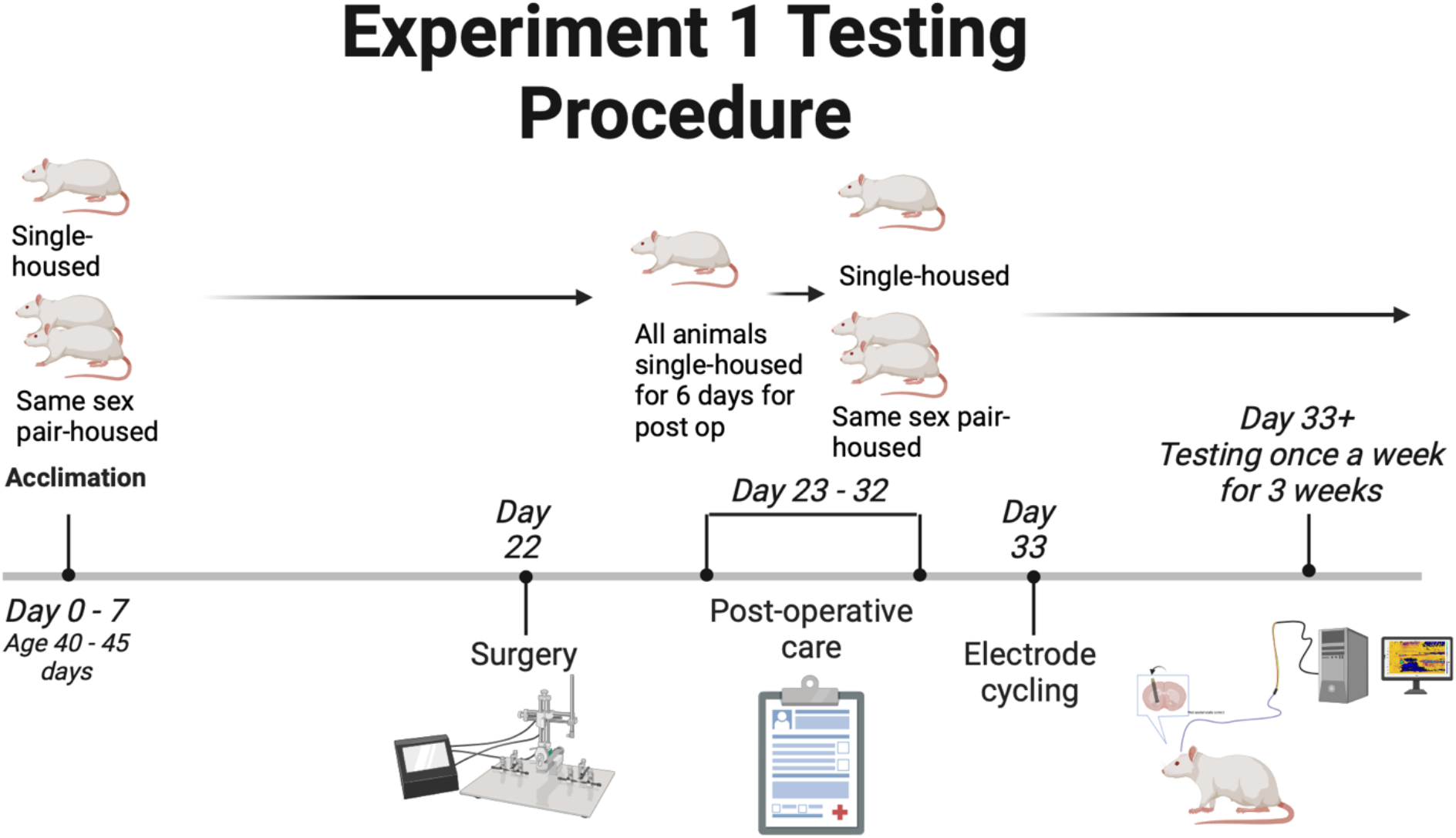
The timeline shows the testing procedure that was followed for experiment 1. It also highlights the social housing situations and when animals were separated. Created with BioRender.com

### Experiment 2

For experiments targeting the DLS of intact rats, 11 male and 11 female rats were used, and they were approximately 45-55 days old on arrival. The animals were kept as social pairs for at least 3 weeks and were separated following surgery until the end of the experiment Figure 2A).

**Figure 2.**
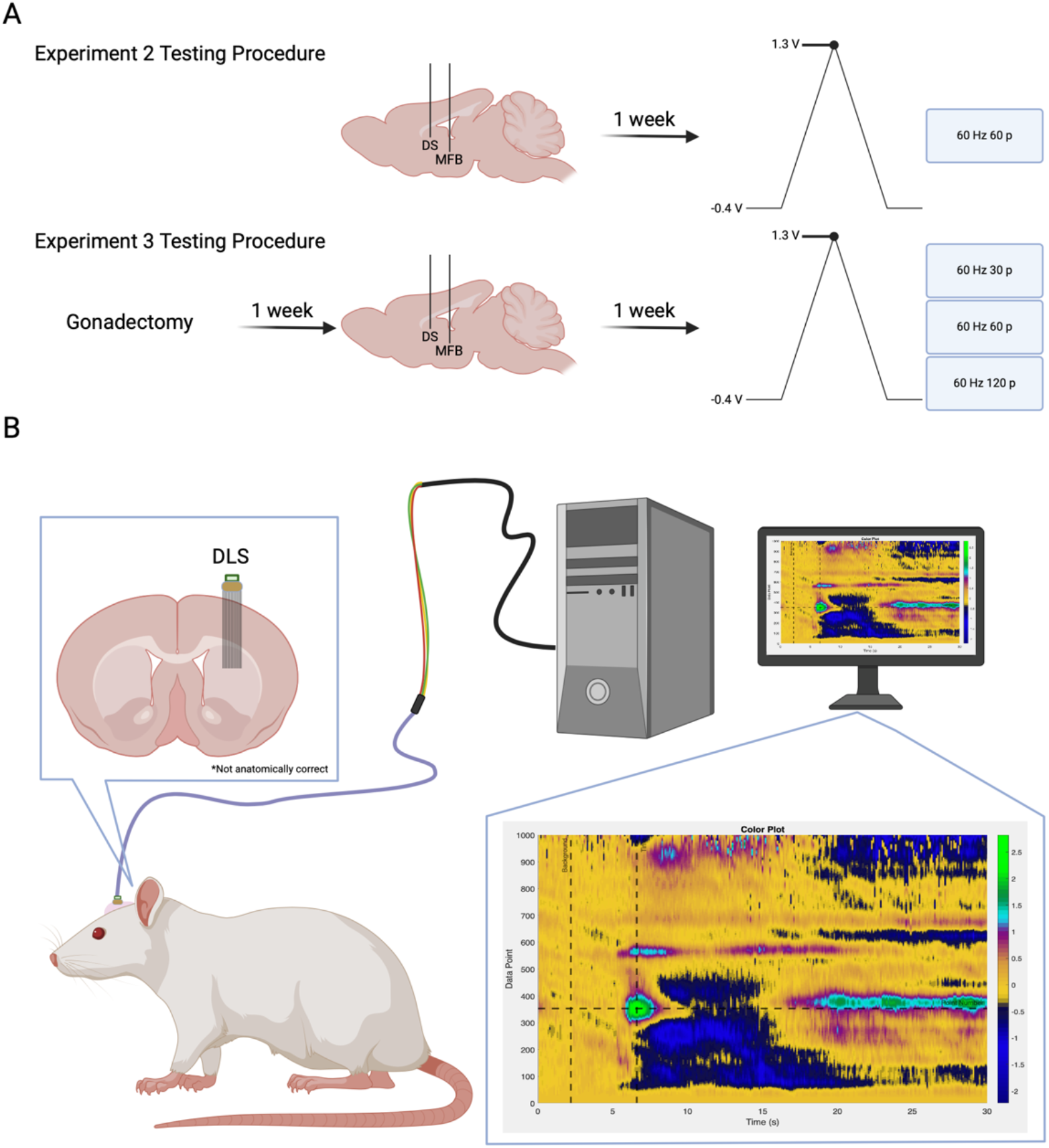
Experimental flow diagram and setup. **A**, Experimental flow and time course for each cohort of animals. **B**, Experimental setup for all animals. Electrodes were either implanted in the NAc or DLS. Animals were connected to the acquisition system via a headstage and cable during behavioral experiments. Created with BioRender.com

### Experiment 3

For experiments targeting the DLS of gonadectomized rats, 5 male and 5 female rats were used, and they were approximately 70-75 days old on arrival. The animals were initially socially housed until gonadectomy surgery and separated following surgery (Figure 2B).

### Acquisition Hardware and Software

Data was acquired using two PCIe-7841 R series cards (National Instruments). Each card acquired eight independent streams of analog data, totaling 16 channels for the system. One of the analog outputs from the first card was used to generate the common ramp signal (−0.4 to 1.3 to -0.4 V) at a rate of 400 V/s (8.5 ms duration), after which there was a 91.5 ms hold period at -0.4 V. This signal was continuously repeated at a rate of 10Hz. If the rate was increased to 60Hz, the software adjusted the hold period accordingly, but the ramp signal stayed the same.

The ramp signal was applied to the positive input of the headstage amplifier (Analog Devices, ADA4062-4ACPZ-R7), while the negative input was connected to the carbon fiber electrode (Takmakov et al., 2011). The amplifier was powered with ± 15 V (B&K Precision, 1672) and had a gain of 200 nA/V with the use of a 4.99 MΩ resistor. The output of each amplifier was fed into one of the analog inputs’ positive terminals on the R series cards (i.e., one fiber per analog input). The ramp signal applied to the amplifiers was also sent directly to the same analog inputs’ corresponding negative terminals. The differential of these two signals is the measured signal.

Connecting the R series cards to the headstage used a variety of adapters and interfaces. Briefly, SHC68-68-RMIO cables (National Instruments) were connected to a custom interface box from the R series cards. The output of this box was a DB25 connector that was interfaced to an Omnetics A79029-001 connector (Omnetics Connector Corporation) using a 24-conductor wire bundle (Digikey, MB24S-50-ND). The A79029-001 connector mated with the headstage, which had connector A79024-001 from Omnetics.

The acquisition software was written in LabVIEW (National Instruments) and acquired 1000 data points during each ramp signal. Each channel had its data saved to separate files in 30-second increments; however, this value can be changed based on experimental needs. If enabled, the software generated a stimulation trigger signal that was routed through another analog output of the first R series card. This trigger signal was then routed to a stimulus isolator (ISO-Flex, A.M.P. Instruments, Jerusalem, Israel) that delivered a stimulation current to an implanted stimulating electrode. The stimulation trigger signal was programmed to be applied during the -0.4 V hold period so as not to interfere with data acquisition during the ramp period.

A complete guide to the system’s hardware and software specifications can be found at https://chestekresearch.engin.umich.edu/carbon-fibers/.

### HDCF Array (Working Electrode) Calibration

To provide calibration data, the 16-channel HDCF array (Huan et al., 2021), functionalized with 50 μm laser-ablated tip sites and treated with plasma ashing (Patel et al., 2020), was lowered into a flowcell that was constructed in-house. Artificial cerebrospinal fluid (aCSF) (125 mM NaCl, 1.3 mM CaCl_2_, 4 mM KCl, 2.66 mM PO_4_, 1 mM KCl_2_) made with 3x filtered water and brought to a pH = 7.2-7.4, was kept at a constant flow rate of 100 mL/hr. A silver | silver chloride (Ag|AgCl) reference electrode provided a chemical reference. The reference electrode was made by submerging one end of a silver wire (Sigma, 265 586) in 2N hydrochloric acid (HCl) (ThermoFisher, SA431500) while the other end was connected to the cathode end of a power supply. A silver wire from the same manufacturer, also submerged in the HCl, was connected to the anode end of the power supply. The power supply was set to 5 V and turned on for 30 sec.

The device was first cycled at a rate of 60Hz for 5-10 min and then 10Hz for 30 min. This ‘cycling session’ stabilizes the working electrode signal. The device was then pulsed with 3 bolus each of 4 different concentrations of dopamine hydrochloride (Sigma, H8502-10G), 0.5 μM, 1 μM, 2 μM, and 4 μM, in aCSF at 250 mL/hr. During the dopamine oxidation period, values were measured three times: at the peak of the onset, just before the falloff from the washout, and then in between these two times. The measurements were then averaged into a calibration curve for the devices per cohort. The fiber tips were dipped and held in 3% hydrogen peroxide for 7 seconds and then isopropyl alcohol for 7 seconds to clean the carbon fiber surface and remove any contamination that had bonded to the carbon (Patel et al., 2020). Devices were sterilized with ethylene oxide.

### Experiment 1, Part A: Fast-Scan Cyclic Voltammetry Surgery in the NAc

12 female and 10 male rats were anesthetized with a combination of ketamine (60 mg/kg, i.p.) and dexmedetomidine hydrochloride (0.3 mg/kg, i.p.). A 3-arm stereotaxic apparatus was used with custom multiangle device holders. The first opening through the skull was made for the Ag|AgCl reference electrode cannula in the contralateral cortex from the working electrode coordinates. A second opening was made in the skull at the site to implant the stimulating electrode. A third opening was made in the skull centered over the working electrode coordinates to form a 2.8 mm x 1.4 mm oval. Next, holes were drilled along the perimeter of the incision site for 6-8 bone screws (Grainger, 1ZA99) to secure the devices to the skull. Screws are hand-driven into place, and then a layer of Metabond dental adhesive (Henry Schein, 1865548) was added, making sure to thoroughly coat each of the skull’s screws while avoiding openings created for the three electrodes. Next, the dura was resected while preserving the cortex. Eye spear surgical sponges absorbed Blood and cerebrospinal fluid (Henry Schein, 1245120).

A reference electrode cannula (BASi, MD-2251) was trimmed 5 mm below the pedestal. Next, the Ag|AgCl wire was glued into the cannula’s mating connector and inserted into the cannula, extending 2mm past the end of the trimmed tube. An acute reference electrode in the cannula was lowered into position (AP -3.0 mm, ML ±3.5 mm, DV (relative to the skull) -1.2 mm). A bipolar stimulating electrode (Plastics One, Inc., MS303/2-B/2 ELEC .008” TW Bend 6.5mm over, 12mm after bend) was lowered from the top of the skull into the ventral tegmental area (VTA) (AP -5.15-5.2 mm, ML ±0.75-0.8 mm, DV (relative to the skull) -8.2-8.4 mm). The working electrode’s fiber tips were brought to the brain surface, and the FSCV system was turned on. The stimulating electrode was lowered 0.2 mm every 10 sec, while the HDCF array was slowly lowered 0.1 mm every 10 sec from zero, which was the top of the brain, into the NAc (AP +1.35-1.4 mm, ML ±2.9-3.2 mm, DV (relative to the cortical surface) -6.8-7.2 mm, at an angle of 16.7 degrees). Unlike more rigid electrodes, the flexible carbon fibers can be deflected by the tough epithelial cells of the lateral ventricles when targeting the medial region of the NAcS. To avoid this problem, the electrodes were implanted at an angle. Using the custom LabVIEW software, a stimulus isolator was used to apply stimulation pulses to the stimulating electrode. These stimulation pulses were used to stimulate DA release for detection by the working electrode once in place.

Electrodes were anchored in place with the use of dental acrylic. The working and stimulating electrodes were covered with protective dust caps (Omnetics, A79041-001 and Plastics One, Inc., 303DCMN), and the reference electrode cannula was occluded with the provided sterile stylet. After the surgery, rats were given atipamezole hydrochloride (0.5 mg/kg i.p.) and dexamethasone (200 μg/kg s.c.). The rats then went through standard post-operative care.

### Experiment 1, Part B: Fast-Scan Cyclic Voltammetry Testing (NAc)

On day 11 post-surgery, animals underwent a cycling session where an Ag|AgCl reference electrode was lowered into the reference electrode cannula, and a headstage was connected to the working electrode. The stimulating electrode was connected to the isolation unit. The reference electrode was connected to the headstage using a gold pin connector. The working electrode was repeatedly cycled at a rate of 60Hz for 10-15 min or until the detected current was stable for 5 min.

The waveform was then cycled at a rate of 10Hz for 20-30 min or until the standard deviation for the detected current was consistently below 0.4 nA for 5 min. This ‘cycling session’ stabilizes the working electrode signal. On testing days, a cycling session occurs before any recordings are taken, and then a 10-minute ES DA recording is collected with the applied waveform repeatedly cycled at a rate of 10Hz. All data recordings were collected at 10Hz. Following the ES DA collection, three 30-second stimulation events recordings were taken, with 5 min between each recording. Five seconds into each recording, a pulse train was applied to the stimulating electrode using a stimulus isolator controlled by custom LabVIEW software. Three stimulation parameters were used for our stimulation events: pulse trains of 30Hz 15 total pulses (p), 60Hz 30p, or 60Hz 60p. During the initial stimulation test session, the peak amplitude of the current applied was optimized for each rat, ranging from 120-200 μA, and held constant for all pulse trains (Example stimulation parameter: 30Hz 15p, 60Hz 30/60p 150 μA).

Animals were monitored during each stimulation trial, and their physical reactions were recorded. Following the last stimulation event, the working and stimulating electrodes were disconnected from their respective cables and hardware, and their dust caps were replaced. The reference electrode was removed and replaced with a sterile stylet.

Vaginal smears were taken every day and before FSCV testing occurred except during surgery recovery days. In the NAcC and NAcS, 8 females were in the metestrus/diestrus phase and 2 in the proestrus stage.

### Experiment 2, Part A: Fast-Scan Cyclic Voltammetry Surgery in the DLS

The FSCV surgery protocol was the same as previously described. Coordinates were modified for the DLS for the working electrode, and the stimulating electrode was implanted into the medial forebrain bundle (MFB). Coordinates in relation to bregma: working electrode targeting the DLS: (AP +0.2 mm, ML ±3.0 mm ± 1.0 mm, DV (relative to the cortical surface) -4.0 mm), bipolar stimulating electrode (Plastics One, Inc.) targeting the MFB: (AP -1.8mm, ML ±2.0 mm ± 0.5 mm, DV (relative to the skull) -7.8 mm for males and -7.5 mm for females), and reference electrode guide cannula in the contralateral cortex (AP -3.0 mm, ML ±3.5 mm, DV (relative to the skull) -1.2 mm).

### Experiment 2, Part B: Fast-Scan Cyclic Voltammetry Testing (DLS)

The cycling session followed the same procedure as described above, and animals were tested during the beginning of their light/dark cycle. The testing days followed a similar procedure as described above. Intact female and male rats were allowed to cycle for 10 min at 10Hz, and then a single electrical stimulation was applied at 60Hz 60p. The animal was given an injection of methamphetamine (1.0 mg/kg s.c.) and then cycled for 10 more minutes before taking another 60Hz 60p stimulation. Peak amplitude was determined in an identical fashion as described above. Vaginal lavage was taken daily before FSCV testing.

### Experiment 3, Part A: Gonadectomy

Females were ovariectomized (OVX), and males were castrated (CAST) under isoflurane anesthesia approximately one week prior to the FSCV surgery. Briefly, OVX was performed via a single dorsal incision along the midline below the ribcage. One incision is made on each side laterally through the muscle wall. Each ovary is externalized and then removed following cauterization of the connective tissue. The tissue is returned to the abdominal cavity, and the muscle is sutured on each side. The external incision is closed with an 11 mm wound clip. For CAST, the testes are removed via a ventral approach. A small incision is made along the bottom of the scrotum. The scrotal sac is opened to externalize each testicle and vas deferens before removal. The connective tissue is sutured with surgical thread. The incision site is closed via an 11 mm wound clip. Animals are given one week to recover. Following recovery, vaginal lavage samples were collected from females to confirm the absence of the estrous cycle. Additional details regarding surgeries are published in Hu and Becker, 2003.

### Experiment 3, Part B: Fast-Scan Cyclic Voltammetry Surgery (DLS)

The FSCV surgery protocol was the same as in Experiment 2.

### Experiment 3, Part C: Fast-Scan Cyclic Voltammetry Testing (DLS)

The cycling session followed the same procedure as described above, and gonadectomized (GDX) rats were tested during the beginning of their light/dark cycle. The testing days followed the same procedures for initial cycling and collection of ES DA recordings as described above but with different stimulation parameters. For our stimulation events, the parameters were as follows: 60Hz 30p or 60Hz 60p. During the initial stimulation test session, the peak amplitude of the current applied was optimized for each rat, ranging from 90-150 μA, and held constant for all pulse trains (Example stimulation parameter: 60Hz 30/60p 150 μA).

### Data Processing

All FSCV data was collected and stored as UTF-8 int16 and analyzed using chemometric analysis with principal component regression (PCR). Each CV was filtered through a 4th order, 2 kHz low pass infinite impulse response (IIR) filter in preprocessing to remove non-biologically relevant noise (Montague et al., 2004). A zero-phase 2nd order, 0.01 Hz high pass Butterworth IIR filter was used on each potential step across time to remove background drift due to the charging current (DeWaele et al., 2017).

PCR was performed using an in-house MATLAB code. A standard training set was constructed based on *in vivo* recordings. The current was converted to concentration using the electrode calibration factor obtained before implantation. If the calculated concentration for a channel fell outside of the mean of all 16 channels ± 2 standard deviations, it was identified as an outlier and removed. The training set consisted of 5 DA, 3 acidic pH, and 2 basic pH voltammograms. Voltammograms were collected from multiple animals while implanted in the DLS or NAc.

### Code Accessibility

Custom MATLAB code is available from the corresponding author upon request.

### Experimental Design and Statistical Analysis

Electrically stimulated (ES) DA release is defined as the difference between pre-stim [DA](nM) and the peak height ES [DA] (nM) within 10 sec of the stimulation pulse in all experiments. All statistical analyses were performed using GraphPad Prism v10.0.

#### Experiment 1: NAc

A one-way ANOVA was conducted with all four groups to look at the differences in ES DA in the NAcC and NAcS of males and females. A two-way ANOVA was conducted to examine the brain region differences in ES DA between NAcC and NAcS among groups. In the case of a significant interaction, a Tukey correction was used for multiple comparisons. The threshold for significance for all statistical tests was set to p < 0.05.

#### Experiment 2: DLS

ES DA and Tau, a measure of DA reuptake (Yorgason, 2011), were analyzed with identical methods. A Geisser-Greenhouse correction was used because sphericity was not assumed. Data was analyzed using a 2-way ANOVA to analyze group means looking at the effect of sex and METH on ES DA release. In the case of a significant interaction, a Bonferroni correction was used for multiple comparisons. The threshold for significance for all statistical tests was set to p < 0.05.

#### Experiment 3: DLS

A Geisser-Greenhouse correction was used because sphericity was not assumed. Because not all fibers had a data point at all time points, data were analyzed by fitting a mixed model. Unfortunately, we could not conduct more detailed analyses of specific fibers across time as not all fibers demonstrated responses above the threshold on all days. We assessed group mean differences in ES DA release within a specific pulse parameter (60Hz, 30p and 60p) across time. In the case of a significant interaction, a Tukey correction was used for multiple comparisons. Sex differences in Tau across time were analyzed by fitting a mixed model. In the case of a significant interaction, a Šídák correction was used for multiple comparisons. The threshold for significance for all statistical tests was set to p < 0.05.

### Histology - Sectioning

In order to maintain the placement of individual electrode fibers, the skull was extracted with the implant intact and sectioned using a slice-in-place technique (Patel et al., 2020). Briefly, animals were euthanized with Fatal Plus (Vortech Pharmaceuticals, VPL 9373) and transcardially perfused with 0.1 M phosphate-buffered saline (PBS) followed by 4% paraformaldehyde (ChemCruz, SC-281692). The animal was then decapitated, and the skull was stripped of skin, ensuring the implant remained cemented to the skull. The skull was then soaked in 0.25 M ethylenediaminetetraacetic acid (Sigma, E5134) in 0.1 M PBS solution at 4 °C with low agitation. The solution was changed every 24 hours for 14 days. The softened skull was then transferred into a 30% sucrose (Sigma, S0389) solution with 0.02% sodium azide (Sigma, S2002) for 48 hours or until sectioning. Before sectioning, skulls were immersed in OCT (ThermoFisher, 4585) and put under a -0.06 MPa vacuum. Skulls were frozen and mounted on a cryostat (ThermoFisher, HM 525NX). 300 μm horizontal sections were taken at -19 °C and stored in 0.1 M PBS with 0.02% sodium azide.

### Histology - Immunohistochemistry

Staining protocols were adapted from Patel et al., 2020. All steps were done under low agitation conditions. Sections were selected for histology by visual verification of carbon fiber tips in tissue with an optical stereo microscope (Amscope SMZK-1TSZ, WF 10x/20). Sections were incubated in 4% paraformaldehyde at 4 °C for 24 hours, then given two 1-hour, 1x PBS washes followed by incubation in Starting Block blocking buffer (ThermoFisher, 37538) with 1.5% Triton X-100 (Sigma, 93443) for 24 hours at room temperature. The sections were then washed in 1x PBS with 0.5% Triton X-100 (PBST) three times, 1 hour per wash, and incubated in a primary antibody cocktail containing mouse anti-NeuN (Millipore, MAB277 lot 3574318, 1:500 dilution), chicken anti-tyrosine hydroxylase (TH) (Abcam, AB76422 lot GR3367897, 1:500 dilution), rabbit anti-calbindin (CB) (Swant, CB-28a lot 9.20, 1:500 dilution) or rabbit anti-μ-opioid receptor (M-OR) (Immunostar, #24216 lot 20290001, 1:500 dilution) in 1.5% Triton X-100, 0.02% sodium azide, and 1% donkey serum for 10 days at 4 °C. Following three more 1 hour washes in 0.5% PBST, sections were incubated with secondary antibodies donkey anti-mouse Alexa 647 (Jackson, 715-605-150, 1:500 dilution), anti-chicken Alexa 488 (Jackson, 703-545-155, 1:500 dilution), anti-rabbit Alexa 546 (Life Technologies, A10040, 1:500 dilution), DAPI (Invitrogen, D1306, 1:2000 dilution), 0.5% Triton X-100, 0.02% sodium azide, and 1% donkey serum for 4 days at 4 °C. Finally sections were washed twice, 2 hours each wash, in 0.5% PBST and then stored in 1x PBS with 0.02% azide.

### Histology - Imaging and Analysis

Sections were imaged on a Zeiss LSM 780 confocal microscope using a 10x objective (Zeiss, C-Apochromat 10x/0.45 W M27). Sections were cleared by immersion in 80% 2,2’-thiodiethanol (TDE) (Sigma, 88561) for 45 minutes to 1 hour and mounted on electrostatic slides with slide spacers containing TDE. The active surface of the working electrodes is a cylinder with a height of ∼50 μm and a diameter of approximately 6.8 μm. A z-stack image containing the full 16 fibers with 8 μm steps was used to locate each electrode tip (Fig. 3). Images were analyzed using the Fiji distribution of ImageJ (Schindelin et al., 2012).

**Figure 3.**
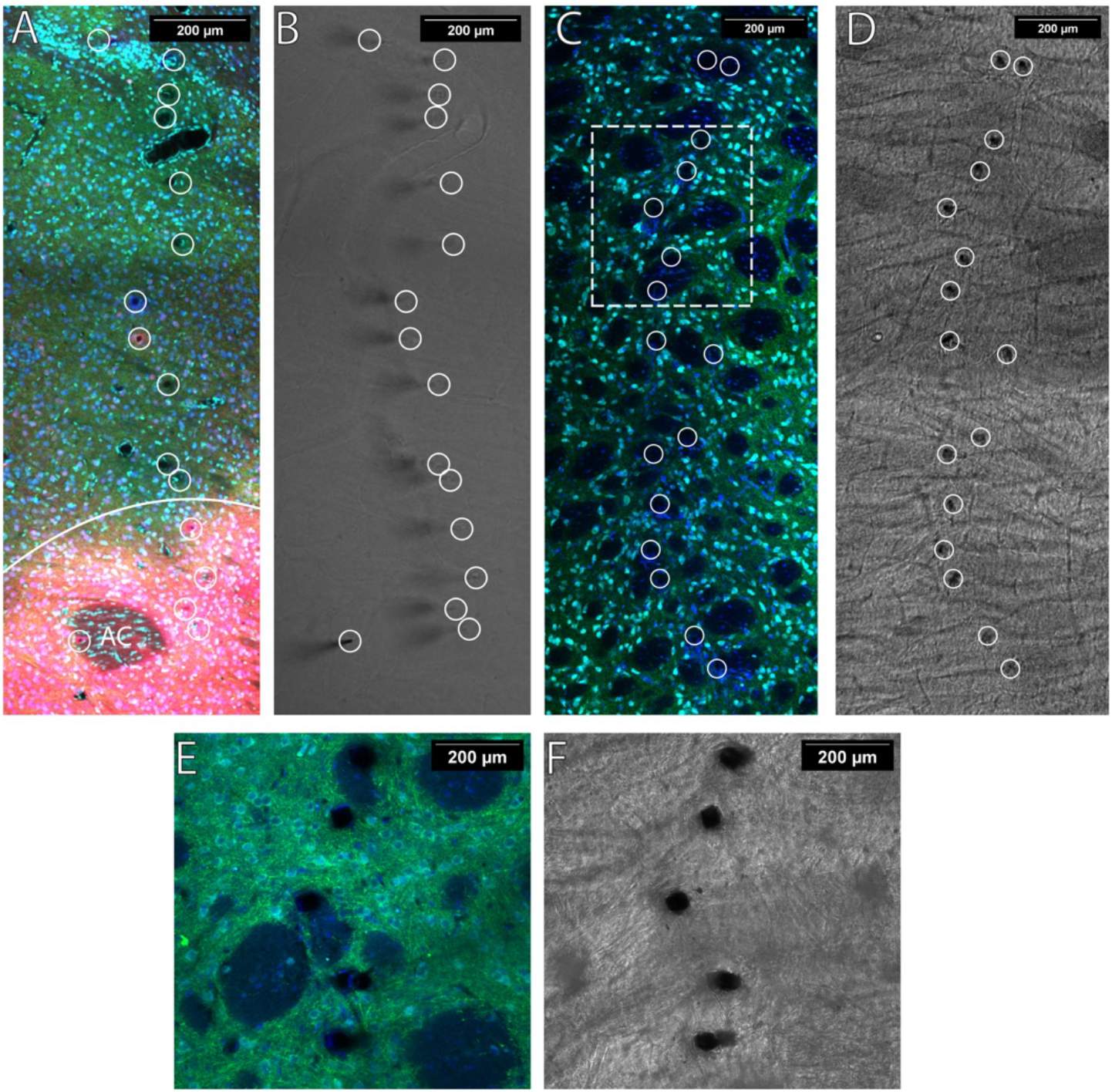
Fiber locations were verified with widefield imaging, and placements are indicated with white circles. TH (green) indicates electrode fibers are placed in DA-containing regions. **A**, When imaging the NAc, Calbindin (red) was used to determine NAcC versus NAcS delineated by the solid white line. The anterior commissure is denoted AC. NeuN (cyan) and DAPI (blue) were used to infer if electrode fibers were placed in a region containing neuronal cell bodies. **B**, Representative widefield image of tissue presented in A. **C, D**, Image with electrode embedded in the DLS with the corresponding widefield image **E**, A close-up image of five representative electrode locations of the same tissue present in panel C. **F**, The corresponding widefield image of E. The contrast of all widefield images was adjusted using the built-in enhance contrast function in Fiji to assist in viewing.

Electrodes targeting the NAc were placed in either the NAcC or NAcS subregions. The NAcC was defined as TH+ and CB+ with strong nuclei and neuropil (Fig. 3A). The NAcS was defined as TH+ and CB- and lateral to the olfactory tubercle or the slight CB+ border region directly flanking the NAcC. The individual electrodes terminate in different DV planes when sectioning extracted tissue horizontally. If more than 8 μm of an electrode was placed in the NAcC or on border regions between the NAcC and NAcS, then the electrode was classified as being located in the NAcC. If more than 24 μm of the active electrode surface was TH- or fell into the olfactory tubercle, they were excluded from the analysis.

A representative image showing TH, NeuN, and DAPI staining in the DLS is illustrated in Figure 3. While not highlighted in this paper, regions of the striatum, such as the DLS, are highly heterogeneous and composed of both patch and matrix compartments (Brimblecombe and Cragg, 2017). These electrodes could also be used to examine differences between the two more closely by comparing the measurements of fibers after verifying their placements using our histological methods.

## Results

### Experiment 1: Nucleus Accumbens - ES DA

A one-way ANOVA was conducted with all four groups to look at the differences in ES DA in the NAcC and NAcS of males and females. When looking at the NAcC at 60Hz 30p, there was a main effect of group (F(3, 159) = 7.455; p < 0.0001; One-way ANOVA; Fig. 4A). In subsequent pairwise comparisons, pair-housed females had significantly higher ES DA release compared to the single females (P<0.0001). Pair-housed females also had higher ES DA release compared to male single-housed males (p = 0.0407) and pair-housed males (p = 0.0154). There were no other differences among the groups.

**Figure 4.**
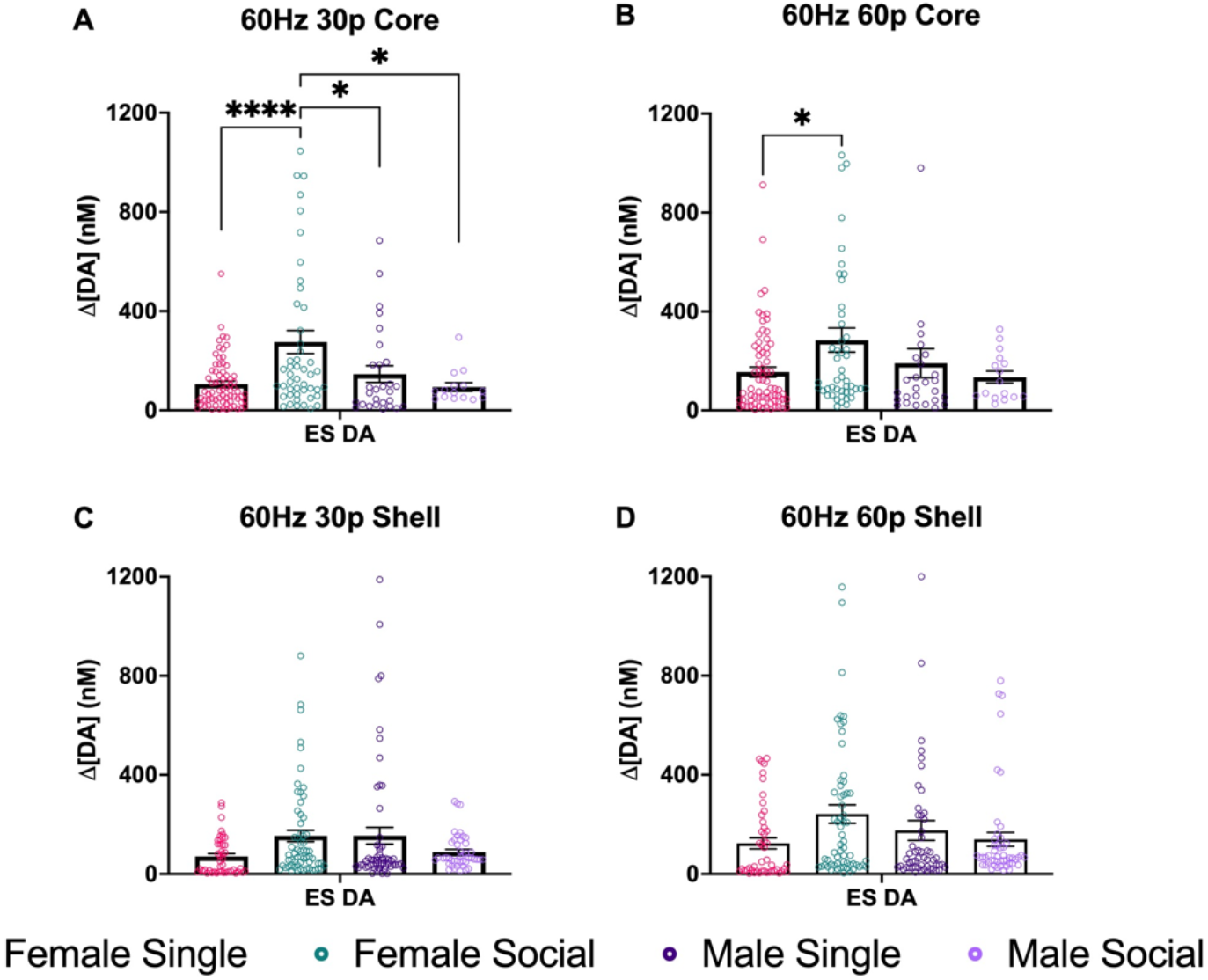
NAc DA release across subregions. **A**, ES DA in the NAcC after stimulation using 60Hz 30p. Here, we found that there was a group effect. Pair-housed females had significantly higher ES DA compared to single females. Pair-housed females had higher ES DA compared to male singles (p = 0.0407) and pair-housed males (p < 0.0154). **B**, Mean max DA in the NAcC after stimulation using 60Hz 60p. There was a significant group effect ((F(3, 158) = 2.946; p = 0.0347)). Pair-housed females had significantly higher ES DA than single females (p < 0.0322). **C**, ES DA in the NAcS after stimulation using 60Hz 30p. No significant effects were found. **D**, ES DA in the NAcS after stimulation using 60Hz 60p. No significant effects were found. Data points indicate values from individual fibers. Bars shown indicate the standard error of the mean (SEM). *p < 0.05, ****p < 0.0001.

At 60Hz 60p, there was also a significant group effect (F(3, 158) = 2.946; p = 0.0347; One-way ANOVA; Fig. 4B). Pair-housed females had significantly more ES DA release compared to the single females (p < 0.0322). In the NAcS, there were no significant differences among groups found at 60Hz 30p (Fig. 4C) and 60Hz 60p (Fig. 4D). A two-way ANOVA was conducted to examine the brain region differences in ES DA between NAcC and NAcS within groups. There was a significant interaction between sex and social housing conditions at 60Hz 30p (F(3, 363p) = 3; p < 0.003; Two-way ANOVA; Fig. 5A). In pairwise comparisons, there was a significant difference between core and shell in socially housed females (p=0.0034). There was no effect of brain region (F (1, 363) = 3.094; p=0.0794; Two-way ANOVA). There were no group differences found at the 60Hz 60p stimulation parameter (Fig. 5B).

**Figure 5.**
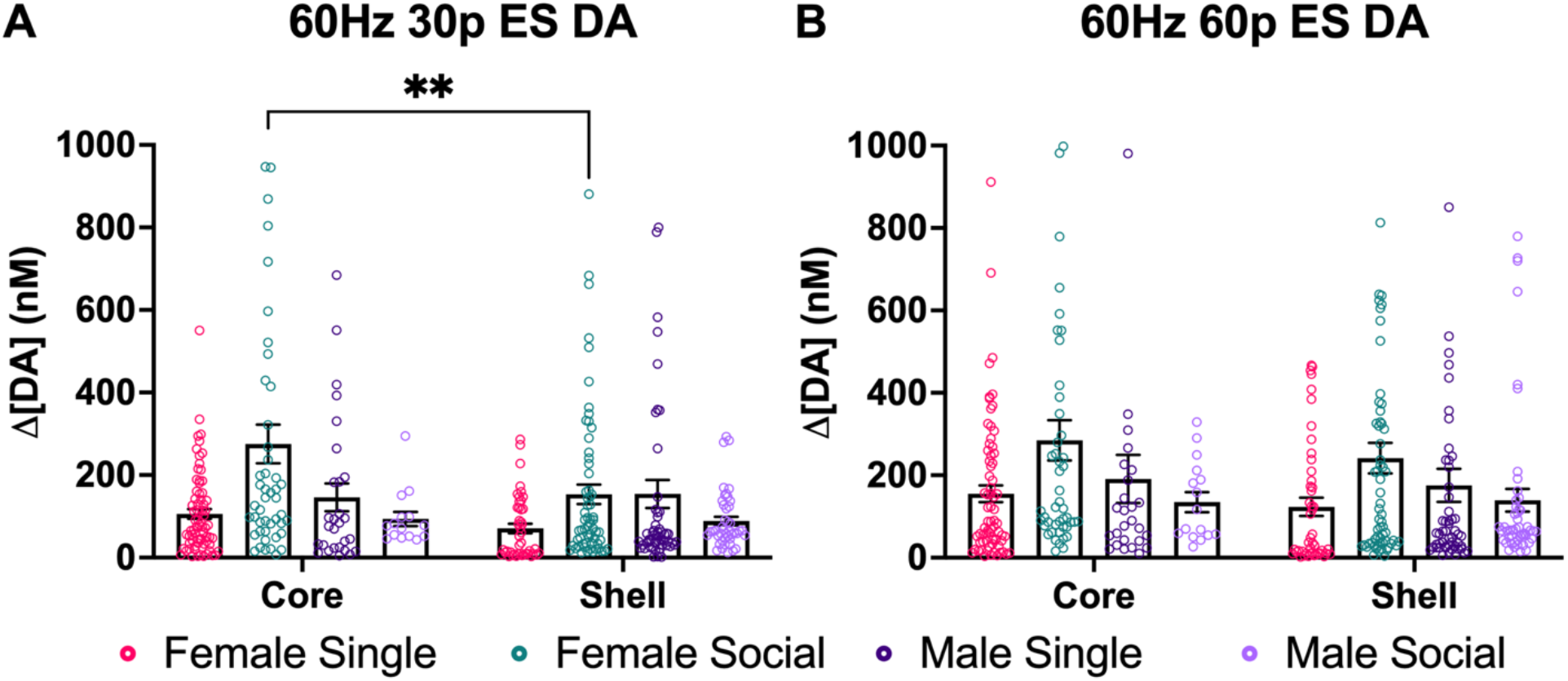
The graphs compare brain regions within groups at two different ES stimulation parameters. **A**, The graphs show differences between NAcC and NAcS at 60Hz 30p. Pair-housed females had higher ES DA release in the NAcC than the NAcS. **B**, No significant differences were found. Data points indicate values from individual fibers. Bars shown indicate SEM. **p < 0.0034.

There was substantial variability among fibers within a brain region and across groups. The coefficient of variation (STDEV/mean) was above 0.5 for all groups and close to 1.0 for many groups. This variability may underlie the failure to find a difference between NAcC vs. NAcS.

When the ES DA response was examined in females in different stages of the estrous cycle, most females were in diestrus, and no differences among stages were found.

### Experiment 2: Sex differences in DLS of gonad-intact animals

ES DA was assessed at a single stimulation parameter before and after administering 1.0 mg/kg METH i.p. in intact female and male rats. There was a significant difference in ES DA release by sex (F(1,617) =15.75; p < 0.0001; 2-way ANOVA; Fig. 6A). There was no effect of METH on the ES DA (F(1,617) = 0.088; p = 0.76; 2-way ANOVA; Fig. 6A) or interaction between METH and sex (F(1,617) = 0.44; p = 0.50; 2-way ANOVA; Fig. 6A).

**Figure 6.**
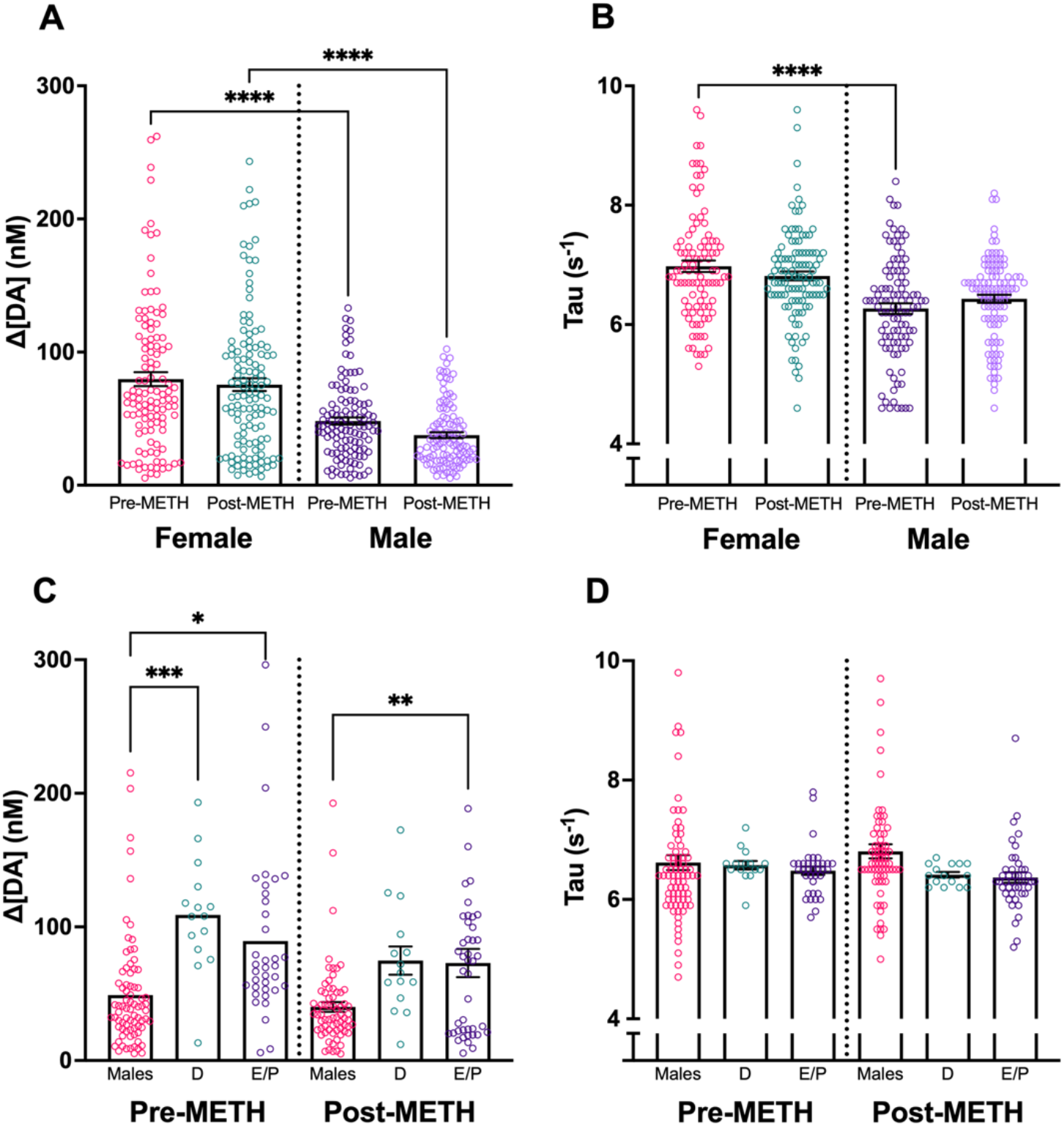
ES DA release before and after i.p. METH injection. **A**, Females exhibited greater ES DA release than males prior to and after METH (1 mg/kg). **B**, Tau was greater in pre-METH females than pre-METH males, but there was no sex difference after METH. **C, D**, ES DA, and Tau comparing males, diestrous females (D), and estrous/proestrus females (E/P). Females exhibited greater DA release than males during both D and E/P phases. There were no differences in Tau among groups. When scaling the figures, three male data points and one female data point were cut off. Data points indicate values from individual fibers. Bars shown indicate SEM. *p < 0.05, **p < 0.01, ***p < 0.001, ****p < 0.0001.

Tau (s^-1^) changes in DA release are a measure of reuptake (Yorgason et al., 2016). There was a significant difference in Tau by sex (F(1,617) = 7.98; p = 0.004; 2-way ANOVA; Fig. 6B), with females having greater Tau than males, indicating slower DA reuptake. There was no effect of METH on Tau (F(1,617) = 0.0036; p = 0.95; 2-way ANOVA; Fig. 6B), or interaction between METH and sex (F(1,617) = 1.79; 2-way ANOVA; p = 0.18; Fig. 6B).

ES DA was also assessed with respect to the estrous cycle stage. There was a main effect of group (F(2,250) = 15.12, p < 0.0001; 2-way ANOVA) with no main effect of METH (F(1,250) = 1.22, p = 0.27; 2-way ANOVA) or interaction between group and METH (F(2,250) = 0.893, p = 0.41; 2-way ANOVA). In subsequent pairwise comparisons (Bonferroni), males (N=6 animals/fibers) had significantly lower ES DA release than diestrous females (N=1 animal/ 16 fibers); p = 0.0001) and estrous/proestrous females (N=3 animals/fibers; p = 0.0004) before METH, there were no differences between estrous cycle stages. Males also had significantly lower ES DA than estrous/proestrous females after METH (p = 0.0047). There were no effects of the estrous cycle on Tau.

### Experiment 3: Repeated electrical stimulation significantly enhances DA release in the DLS in both males and females

ES DA was assessed at two stimulation parameters across four weeks. Stimulation parameters, as well as the timing between stimulations, were picked to minimize carry-over effects on DA release between recordings.

In CAST males, there was a significant effect of week (F(2.116, 182.0) = 45.41; p < 0.0001; 2-way mixed-effects; Fig. 7A) and pulse number (F(1, 127) = 9.259; p = 0.0028; 2-way mixed-effects; Fig. 7A) but no interaction between pulse number and week (F(3, 258) = 1.855; p = 0.1377; 2-way mixed-effects; Fig. 7A). Multiple comparisons revealed a significant difference between week 1 and weeks 2 (p < 0.0001), 3 (p < 0.0001), and 4 (p < 0.0001) as well as weeks 2 and 3 (p = 0.0026) at the 30 pulse parameter. At the 60 pulse parameter, there were significant differences between week 1 and weeks 2 (p < 0.0001), 3 (p < 0.0001), and 4 (p < 0.0001), and between weeks 2 and 3 (p < 0.0127). These results indicate the males’ sensitization as ES DA release increased until week 3, when maximal changes in ES DA release stabilized.

**Figure 7.**
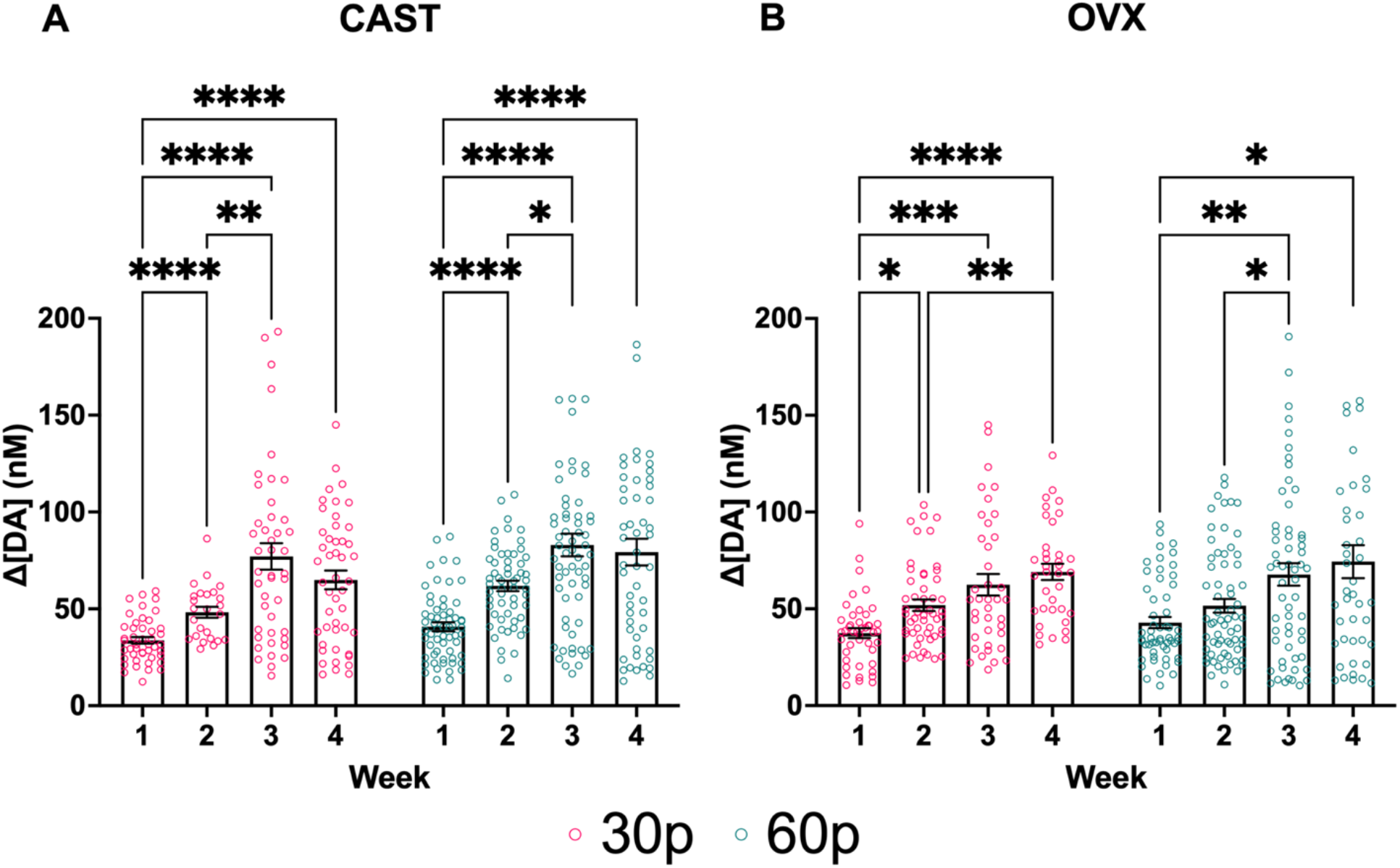
ES DA release increases across time in both **A**, CAST males and **B**, OVX females. Data points indicate values from individual fibers. 3 data points are outside the axis limits for the males, and 2 data points are outside the axis limits for the females. Bars shown indicate SEM. *p < 0.05, **p < 0.01, ***p < 0.001, ****p < 0.0001.

In OVX females, there was a significant effect of week (F(2.393, 195.4) = 15.98; p < 0.0001; 2-way mixed-effects; Fig. 7B), but no effect of pulse (F(1, 140) = 1.003; p = 0.3183; 2-way mixed-effects; Fig. 7B) or interaction between pulse number and week (F(3, 245) = 0.2019; p = 0.8950; 2-way mixed-effects; Fig. 7B). Multiple comparisons at the lowest stimulation parameter revealed a significant difference between weeks 1 and weeks 2 (p = 0.0180), 3 (p = 0.0009), and 4 (p < 0.0001), as well as a difference between weeks 2 and 4 (p = 0.0053). At the 60 pulse parameter, multiple comparisons revealed significant differences between weeks 1 and weeks 3 (p = 0.0072) and 4 (p = 0.0156) as well as between weeks 2 and 3 (p = 0.0442). There were no sex differences in stimulated DA release at any stimulation parameter.

ES changes in Tau (s^-1^), a measure of DA reuptake, were assessed at two stimulation parameters across four weeks (Fig. 8). Stimulation parameters, as well as the timing between stimulations, were picked to minimize carry-over effects on Tau between recordings. At the lowest stimulation parameter, there was a significant effect of sex (F(1, 119) = 7.758; p = 0.0062; 2-way mixed-effects; Fig. 8A) and week (F(1.197, 86.16) = 4.171; p = 0.0373; 2-way mixed-effects; Fig. 8A) as well as a significant interaction between sex and week (F(3, 216) = 5.676; p = 0.0009; 2-way mixed-effects; Fig. 8A). Multiple comparisons revealed significant differences between males and females during weeks 2 (p = 0.0086), 3 (p = 0.0074), and 4 (p = 0.0247). Interestingly, at 30p, female Tau values were lower during week 1 compared to weeks 2-4, indicating faster reuptake at the beginning of the study, as opposed to males who showed no significant difference in Tau at 30p across all four weeks. The higher Tau values of females indicate reduced dopamine reuptake. At the 60Hz 60p stimulation parameter, there was a significant effect of sex (F(1, 150) = 15.63; p = 0.0001; 2-way mixed-effects; Fig. 8B), and week (F(2.188, 228.3) = 7.698; p = 0.0004; 2-way mixed-effects; Fig. 8B) as well as a significant interaction between sex and week (F(3, 313) = 5.947; p = 0.0006; 2-way mixed-effects; Fig 8B). Multiple comparisons revealed significant differences between males and females during week 1 (p = 0.0007).

**Figure 8.**
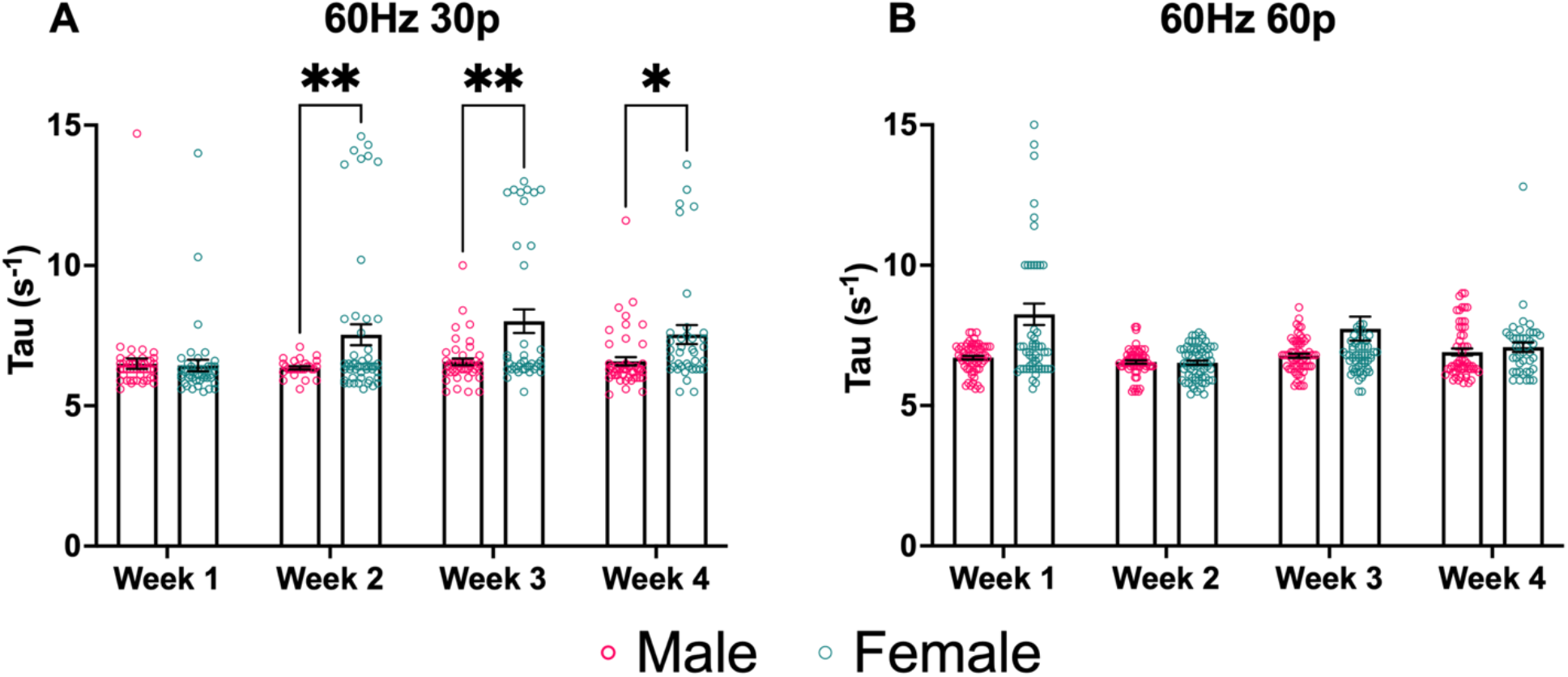
Sex differences in ES dopamine reuptake in males and females are reflected using Tau (s^-1^), a measure of DA reuptake. **A**, At 30p, females had higher Tau values than males during weeks 2 and 3, indicating slower dopamine reuptake. **B**, At 60p, females had higher Tau values than males during the initial test session. Data points indicate values from individual fibers. Three data points from week 1’s female data are outside the axis limits. Five data points from week 3 female data are outside the axis limits. Bars shown indicate SEM. *p < 0.05, **p < 0.01, ***p < 0.001, ****p < 0.0001.

Color plots of DA release from each of the 16 channels of one rat over 4 weeks are illustrated in Figure 9.

**Figure 9.**
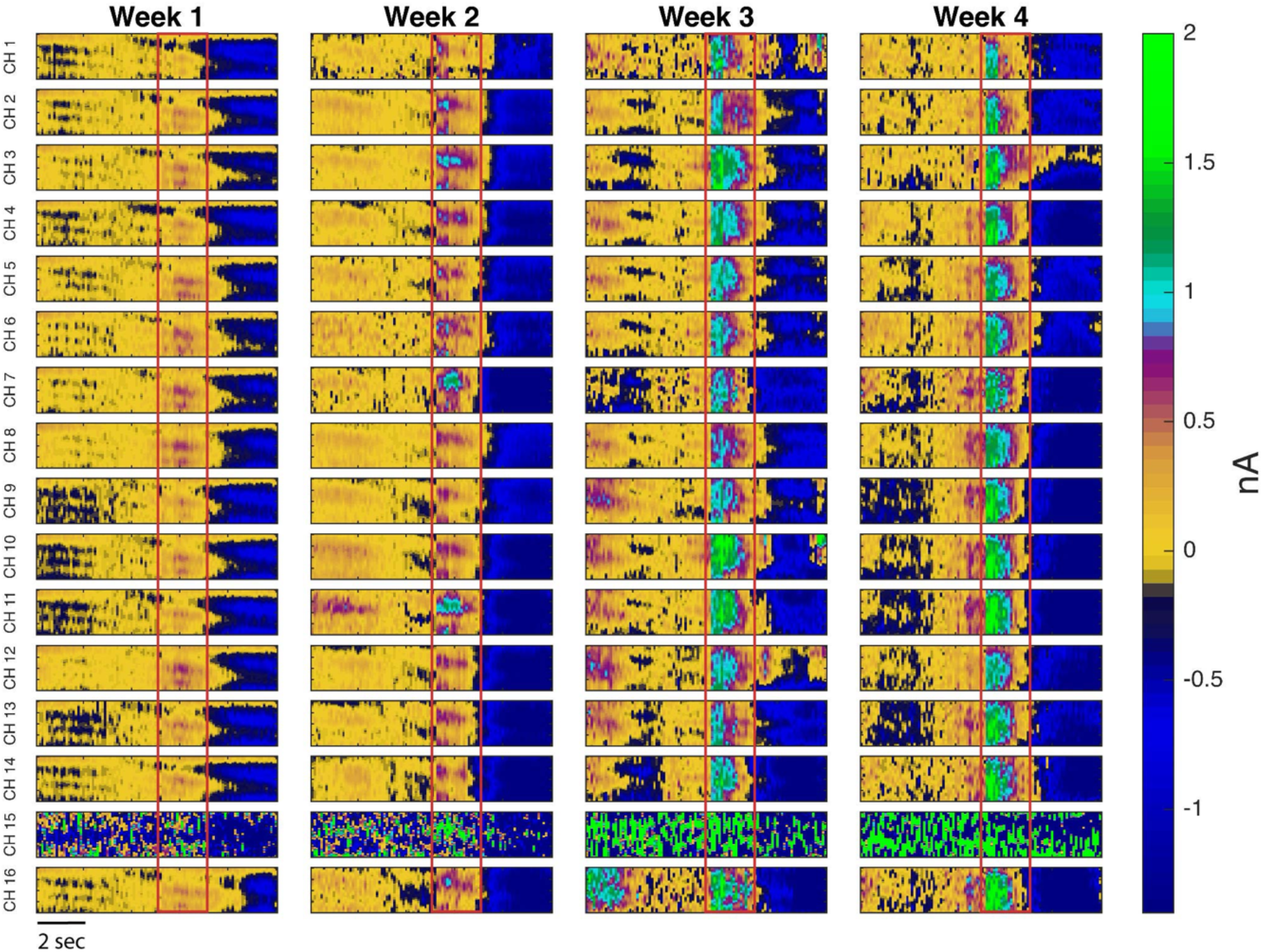
Representative pseudocolor plots of DA release in the DLS of one male rat over four weeks. The red box denotes the onset and conclusion of a 60Hz 30p stimulation train. As can be seen from this color plot, the Δ[DA] (nM) response increased over four weeks. In this particular animal, channel 15 malfunctioned, but channels 14 and 16 functioned as expected.

Table 1 depicts the coefficients of variance (mean ± SEM) for both sexes at the two stimulation parameters. The data show a trend of increasing variability among the fibers over time.

**Table 1.** Sex-specific Fiber Variability: Coefficients of Variance Across Four Weeks.

|  |  |  |  |  |
| --- | --- | --- | --- | --- |
| 30p CV | Week 1 | Week 2 | Week 3 | Week 4 |
| Male | 0.20±0.04 | 0.27±0.08 | 0.30±0.03 | 0.46±0.14 |
| Female | 0.27±0.02 | 0.37±0.07 | 0.49±0.07 | 0.45±0.05 |
| 60p CV | Week 1 | Week 2 | Week 3 | Week 4 |
| Male | 0.28±0.05 | 0.26±0.06 | 0.39±0.09 | 0.51±0.17 |
| Female | 0.38±0.02 | 0.38±0.6 | 0.49±0.08 | 0.40±0.06 |

## Discussion

The NAc consists of the NAcC and the NAcS, each with a different purpose in reward-related behaviors. The NAcC has been associated with wanting, and NAcS has been associated with liking (Morales and Berridge, 2021). Our results show for the first time that housing conditions can differentially affect stimulated DA release, recorded simultaneously in the NAcC vs. NacS, from female rats housed either in pairs or singly. Housing conditions did not affect males’ stimulated DA release in the NAc. These data show us that before any manipulation is introduced, other than housing conditions, there are differences in the VTA DA responsiveness to stimulation in projections to the NAcC that depend on the social housing conditions of the females.

In vitro, phasic DA release in the ventral striatum has been reported to vary between NAcC and NAcS in female mice (Brundage et al., 2022). Here, we found that for the NAcC pair-housed females had higher ES DA release than single-housed females at 60Hz 30p or 60Hz 60p and compared to single-housed males or pair-housed males at 60Hz 30p, but there were no other core-shell differences. Previous findings from the Becker laboratory, using *in vivo* microdialysis, have found that singly-housed females had a greater methamphetamine-induced increase in DA from the NAc (core and shell) compared with pair-housed females (Westenbroek et al., 2019). Differences in the methods for measuring and stimulating DA release certainly contribute to the differences in the direction of the findings. FSCV measures rapid dynamic changes in DA release and isolates NAcC vs. NAcS, while microdialysis sums effects on DA release, reuptake, and diffusion over ∼5-10 minutes. Nevertheless, these studies demonstrate that housing conditions in female rats affect dynamic DA responses to stimulation. Further studies should be conducted to better understand the differences between the subregions of the NAc.

Social isolation in male rats does not affect basal NAc DA in microdialysis studies (Novoa et al., 2021), which is consistent with our findings. Social isolation increased ES-induced DA release in NAc, using FSCV, in adult male mice but did not find an effect of housing in adolescent males (McWain et al., 2022). Another study examined social isolation compared to group housing in males and found that socially isolated males had higher DA release (Yorgason et al., 2016). However, Yorgason et al. (2016) socially isolated their males at twenty-eight days. In our studies, males were pair-housed or isolated beginning at forty to forty-five days of age, and FSCV recording occurred in adults. This suggests that the timing of social isolation is important for effects on NAc DA.

While most studies have only looked at male subjects, a study that looked at male and female copying responses to stress found that social housing positively affected females but not males (Westenbroek et al., 2003). Previous studies also showed that social housing attenuates drug self-administration in females but not in males (Westenbroek et al., 2013; Lee et al., 2017; Moench and Logrip, 2021).

In the DLS of intact animals, we found a sex difference in ES DA before and after methamphetamine was given. In the DLS, we also saw that Tau was higher in females than males; together, this demonstrates greater DA release and lower DA reuptake in females compared to males. These data also indicate a more dynamic DA response in the DLS of females. Previous studies have found that in the DLS of gonad-intact females, the DA active transporter (DAT) is more active and present in higher concentrations than in males (Rivest et al., 1995; Walker et al., 2000; Dluzen and McDermott, 2008). Across several minutes, DA concentrations may appear relatively lower in females than males due to higher reuptake rates. In the studies reported here, we are measuring ES DA release rather than basal activity or activity in slices, which may explain the differences in results reported here. Studies to investigate the effects of ovarian hormones on Tau and stimulated DA in intact and gonadectomized animals are in progress.

Previous experiments from the Becker lab have shown sex differences in basal DA in the dorsal striatum of gonadectomized male and female rats, with CAST males having significantly greater basal DA (Castner et al., 1993; Xiao and Becker, 1994; Cummings et al., 2014; Quigley et al., 2021). However, it is essential to note that these prior studies were conducted using microdialysis, which has a lower temporal resolution, ∼5-10 minutes, compared to the sub-second resolution of the carbon fiber electrodes used here. FSCV measures ES DA increases relative to background rather than concentrations of DA in dialysate, which could be attributed to differences in basal DA release, reuptake, diffusion, or a combination.

Estrous cycle dependent differences in amphetamine-stimulated striatal DA release have been found using microdialysis (Becker and Cha, 1989). FSCV studies have also found estrous cycle dependent differences in striatal DA release (Calipari et al., 2017). Calipari et al., 2017 reported that ES of the NAc elicits greater responses in estrous females than in other stages of the cycle or males. In addition, VTA neuron burst firing rates showed the same elevation pattern in estrous females (Calipari et al., 2017). There was no effect of the estrous cycle in the NAc or in the DLS in females in this study, likely due to the small number of females at different stages of the cycle.

In the DLS, our results showed that stimulated DA release increased from week one to week four at both the 60Hz 30p and 60p stimulation parameters in both gonadectomized males and females, providing evidence for *in vivo* sensitization across time. Amphetamine administration has been shown to induce time-dependent enhancement of DA transmission in the dorsal striatum, compared to saline pretreatment using microdialysis (Paulson and Robinson, 1995). These were one-time measurements after either 3, 7, or 28 days of withdrawal from amphetamine. Only animals tested after 28 days of withdrawal showed a significant enhancement in amphetamine-induced DA release, compared with saline pretreatment, as measured using in-vivo microdialysis (Paulson and Robinson, 1995).

With this novel 16-channel electrode, sensitization of stimulated DA release can be measured in the DLS over 4 consecutive weeks in both CAST males and OVX females. With this chronically implanted electrode, we looked at population differences in stimulated DA release across weeks in the DLS. In contrast to more traditional chronic electrodes, our microarray affords exceptional robustness, which is especially important for chronic electrode preparations. Importantly, our 16-channel device provides redundancy, allowing uninterrupted data collection for extended periods. While, occasionally, DA release fails to be detectable above baseline for a particular electrode, an electrode can come back online during subsequent recordings. Future studies will more directly assess any discernible patterns that appear when assessing fiber functionality over time.

Each electrode on our device functions independent of adjacent electrodes, as proven by differences in responding among channels. This is crucial in depicting the heterogeneity within structures such as the NAc and DLS. Electrode independence, combined with repeated recording sessions, gives our data impressive dimensionality. There are ongoing efforts to increase the performance of our device to minimize the number of channels that fail threshold tests and to increase the signal-to-noise ratio of successful collections, therefore increasing our data yield.

DA signaling within and between these striatal regions has been consistently tied to learning, motivated behaviors, and addiction vulnerability. Differences between males and females are also believed to be a primary driver for sex differences in the escalation of addiction-like behavior (Quigley et al., 2021). However, reliably assessing DA release in these regions in freely moving subjects over time has been difficult. The unique combination of high temporal and spatial resolution provided by these novel 16-channel chronic FSCV electrodes opens up numerous exciting avenues for future research. Altogether, these data support an exciting new method for studying differences in neuronal activity across time as we can better study the role of these highly dynamic and heterogeneous populations of neurons in motivated behavior and any sex differences therein.

In conclusion, our findings demonstrate that in female rats, social housing conditions enhance ES DA release in the NAcC. This same effect of housing is not seen in the NAcS or in males in either subregion of the NAc. In the DLS, we show that intact females exhibit greater ES DA release than intact males before and after methamphetamine. The data indicate that there are sex differences that significantly affect DA reuptake in the DLS, with females having lower DA reuptake levels than males in intact and gonadectomized rats. Furthermore, repeated ES causes sensitization of DA in the DLS of both gonadectomized males and females, as indicated by enhanced ES DA release across weeks.

Finally, these results demonstrate the viability of these novel electrodes to chronically detect DA release from multiple locations in subregions of the striatum of freely moving rats. We successfully monitored DA release in the NAcC and NAcS simultaneously with a single device and measured DA release in the DLS over 4 weeks.

## Acknowledgments

These experiments were supported by grants to JBB from the USPHS: NIH R01DA049795 and R01DA046403 to JBB, T32-DA-007281-26 to S. Flagel & JBB, and UF1NS107659 to CAC

## Notes

Conflict of interest statement: The authors declare no conflicts of interest.

### Competing Interest Statement

The authors have declared no competing interest.

### Summary of Updates

The manuscript has been revised to include additional experiments. The Introduction and Discussion have been revised to provide a more thorough background and to better integrate the experiments.

